# ARF-1 Coordinates Cargo Sorting at FERARI Endosomal Recycling Hubs

**DOI:** 10.64898/2026.01.12.698965

**Authors:** Jachen A. Solinger, Curdin A. Franziscus, Trian Nuredini, Dominic L. Müller, Petia Adarska, Francesca Bottanelli, Anne Spang

**Affiliations:** Biozentrum, University of Basel, Spitalstrasse 41, CH-4056 Basel, Switzerland; Freie Universität Berlin, Institute for Chemistry and Biochemistry, Thielallee 63, D-14195 Berlin, Germany

**Keywords:** tether, recycling endosomes, endocytosis, *C. elegans*, small GTPases, Sec1/Munc18 protein, SNAREs, vesicle transport, Rab proteins, mammalian cells, CRISPR, membrane fusion, membrane fission, adaptor proteins, sorting nexins, *S*. *cerevisiae*

## Abstract

Endosomal pathways are central to cellular organization, compartmentalization, and intercellular communication. Independent of the endocytic entry route, sorting endosomes function as critical junctions at which membrane proteins are sorted for recycling or degradation. The multicomponent tether FERARI orchestrates recycling by coordinating Rab GTPase activity with vesicle fusion and fission through a kiss-and-run mechanism. Here, we identify the small GTPase ARF-1 as an essential regulator of cargo loading at FERARI-dependent endosomal kiss-and-run (KAR) sites. Cycling of ARF-1 between its active and inactive states is required for efficient cargo capture into recycling vesicles. Our findings support a model in which cycling ARF-1 activity governs the temporal association of cargo with adaptor proteins and complexes, thereby controlling selective uptake into vesicles destined for the plasma membrane. These results uncover a previously unappreciated layer of regulation of endosomal recycling and highlight the interplay between ARF and Rab family GTPases in this process.

## Introduction

Efficient mechanisms of uptake and recycling of proteins and solutes are paramount for cells to interact with their environment. They need to exchange materials and signals by a process called endocytosis. In this process, cargo is internalized into cells and reaches endosomes. From there, the cargo can either recycle back to the plasma membrane or to the Golgi/trans-Golgi network (TGN) to ultimately reach the cell surface. Cargo, which is not recycled, will remain in endosomes and ultimately degraded in endolysosomes. While endocytosis is well-studied, the recycling and sorting to the plasma or the TGN remains still less clear (Cullen and Steinberg, 2018; Gopaldass et al., 2024; Scott et al., 2014; Simonetti et al., 2023; Spang, 2016). One reason for this is that there are multiple pathways, which are at least partially overlapping and that recycling can happen from early, from maturing and from late endosomes (Cullen and Steinberg, 2018; Gopaldass et al., 2024; Podinovskaia and Spang, 2018). Some of the pathways returning cargo to the plasma membrane involve a complicated network of dynamic membrane tubules called the tubular endosomal network (TEN) (Grant and Donaldson, 2009; Klumperman and Raposo, 2014). Most research has focused on the initial steps of protein sorting and recycling, which involve tubular structures emanating directly from Rab5-positive early endosomes (Gallon and Cullen, 2015). These transport pathways use complexes such as retromer and retriever and can sort many different cargoes to different destinations (Courtellemont et al., 2022; Cullen and Steinberg, 2018; Gopaldass et al., 2024; Simonetti et al., 2023). The tubular membranes are formed at the sorting endosome and stabilized by sorting nexins (SNXs) which lead to the recruitment of cargo into these tubes by cargo adaptors. Transport carrier vesicles or tubules positive for Rab GTPases such as Rab11 pinch off from the sorting tubules (Chen et al., 2019). It was largely assumed that those carriers would proceed directly to the plasma membrane or the TGN. While this might be the case for cargoes transported to the TGN, it remains less clear, whether this direct transport also occurs for all cargoes destined to the plasma membrane. The large TEN, which is consistently observed in electron micrographs cannot easily be reconciled with the short linear tubules postulated by the direct transport model (Franke et al., 2019; Klumperman and Raposo, 2014; Murk et al., 2003; van der Beek et al., 2022). Also, the low-affinity interactions between cargoes and adaptor proteins (Ghai et al., 2011; Honing et al., 1997; McMillan et al., 2021; Solinger and Spang, 2022; Yong et al., 2020) is very error prone and does not explain the apparent specificity of transport in one single step. An iterative process of sorting would reduce the level of erroneous sorting. Recently, we discovered that RAB-11 and RAB-10 recycling carriers and incoming RAB-5 endocytic carriers undergo kiss-and-run at sorting endosomes and TEN and that during this process cargo was exchanged between the carriers and the sorting endosomes and TEN (Solinger et al., 2020; Solinger et al., 2022). Importantly, the kiss-and-run process was strictly dependent on the FERARI tethering complex. FERARI is a cousin of the HOPS and CORVET tethering complexes, which act in the endolysosomal/autophagosomal pathways and in homotypic fusion of Rab5 compartments, respectively (Shvarev et al., 2024; Solinger and Spang, 2013; Solinger and Spang, 2014; Ungermann and Moeller, 2025), and belong to the CATCHR family of tethers (Chou et al., 2016; Szentgyorgyi and Spang, 2023) (Fig. 1a). Like HOPS and CORVET, FERARI contains Rab interacting subunits as well as an Sec1/Munc13 family (SM) protein for SNARE interaction. In contrast to other known tethers, FERARI also comprises a pinchase activity, which is essential for the fission during the kiss-and-run process (Deo et al., 2018; Solinger et al., 2020) (Fig. 1a). Based on our previous work, we proposed a model in which the sorting endosome/TEN would be used as a platform for protein sorting, where different recycling vesicles could deposit or retrieve cargo, while cargo that was just endocytosed, including potential bystander cargo, could be recycled early on through compartments close to the plasma membrane. By sequential kiss-and-run events a gradual enrichment of cargo would be achieved, similar to a distillation process, making the sorting process more robust and faithful and overcoming missorting due to low-affinity binding between cargo and cargo adaptors (Solinger et al., 2022; Solinger and Spang, 2022).

**Figure 1:**
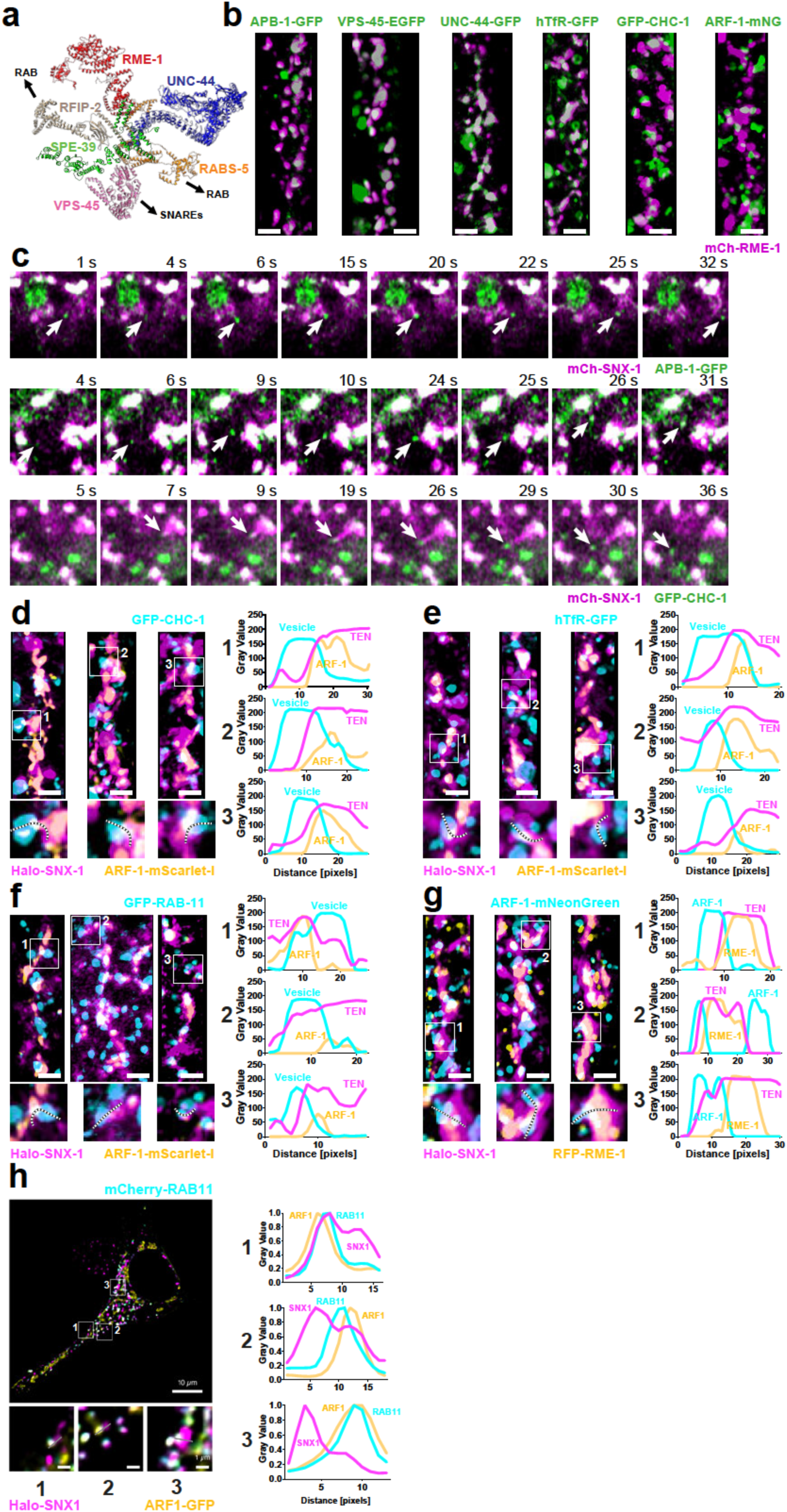
FERARI forms exit sites on SNX-1 compartments that contain ARF-1, cargo and cargo adaptors. (a) Schematic representation of FERARI machinery, showing single proteins folded in Alphafold, arranged according to known interactions. (b) Close association of hTfR, VPS-45, UNC-44, APB-1 (probably AP-1), ARF-1 and CHC-1 (clathrin) with RME-1. 3D projections of z-stacks with 2 µm scale bars. These data indicate the presence of KAR sites with FERARI. (c) Movie stills showing kiss-and-run of APB-1 vesicle (top) and kiss-and-run of CHC-1 (clathrin) vesicle (middle). Formation of a CHC-1 vesicle from a tubule emanating from SNX-1 compartment (bottom). Arrows point at respective vesicles. (d-f) 3D projections showing close juxtaposition of vesicles with SNX-1 compartments containing ARF-1 nanodomains. Vesicles are often in close proximity to ARF-1 domains. Enlargements show selected vesicles with corresponding line plots (on the right) along the vesicle and turning into the TEN to illustrate that ARF-1 is often found between vesicles and SNX-1 compartments. (d) Analyses with CHC-1 vesicles (e) with hTfR vesicles and (f) with RAB-11 vesicles. (g) ARF-1 and RME-1 are juxtaposed on SNX-1 compartments. Scale bars are 2 µm. (h) Rab11 vesicles containing ARF1 can be observed in close proximity to SNX1 compartments in HeLa cells. Enlargements show three examples with line plots demonstrating the overlap between Rab11 and ARF1 as well as the close association with SNX1 (graphs on the right).

We have shown previously that cargo can be transferred from RAB-5 carriers into sorting endosomes/TEN and from there into RAB-11 carriers (Solinger et al., 2022). Still, the directionality remained unclear and we suggested that differential affinities between cargo and cargo receptor might provide directionality. In agreement with our model, it was shown previously that AP-1 and AP-2 cargo adaptor complexes exist in open and closed conformations, in which the open conformation would bind to cargo, while the closed conformation would not (Beacham et al., 2019; Ren et al., 2013). The switch between the two conformation is regulated by the small GTPase ARF-1 (Beacham et al., 2019; Ren et al., 2013).

Here, we provide evidence that ARF-1 cycling between the GDP- and GTP-bound states is required for efficient cargo loading into RAB-11 carriers at sorting endosomes/TEN in *C. elegans*. The cycling between the two states at the sorting endosome/TEN is regulated by the guanine nucleotide exchange factors (ArfGEFs) GBF-1 and AGEF-1 and the GTPase-activating proteins (ArfGAPs) RFGP-1 and RFGP-2, which are the homologues of GBF1 and BIG1/2 and ArfGAP1/2 in mammalian cells, respectively. We suggest that ARF-1 activity cycles regulate the release and re-binding of cargo by cargo adaptors and/or adaptor complexes. This process subsequently drives the enrichment of cargo into RAB-11 carriers in a process, which is evolutionary conserved. We propose that the spatiotemporally restricted re-binding mechanism offers a plausible mechanism ensuring directional cargo sorting through the fusion pore during kiss-and-run. Moreover, our data reveal a dual requirement of ARF-1 in recycling through the TEN. First, ARF-1 activated by GBF-1 is essential for the proper formation of the TEN and second ARF-1 activated by AGEF-1 is required for efficient sorting at the TEN.

## Results

### ARF-1 is present at FERARI docking sites during endosomal recycling

We had previously shown that AP-1 vesicles undergo kiss-and-run on SNX1 positive compartments in mammalian cells and that knock-down of AP-1 subunits caused an enlargement of SNX-1 structures in the *C. elegans* intestine (Solinger et al., 2022). We assumed that this kiss-and-run process is FERARI-dependent and that cargo would be present at these endosomal KAR (*K*iss-*A*nd-*R*un) sites (Solinger et al., 2022; Solinger and Spang, 2022). If this assumption was correct, we would expect that AP-1 should co-localize with FERARI subunits. To this end, we determined the localization of the ß-subunit of AP-1 (APB-1) with the FERARI subunit RME-1 (Fig. 1b). Since FERARI subunits have also other functions in the cell, we first demonstrated that two other FERARI subunits, VPS-45 and UNC-44 are found on the same structures (Fig. 1b, Fig. S1a). As predicted, the cargo hTfR-GFP localized to these sites. We found that APB-1 is present on or juxtaposed to FERARI-positive structures. AP-1 is part of a subset of clathrin-coated vesicles. As expected, clathrin heavy chain (CHC-1) showed a similar localization (Fig. 1b). Consistent with our previous data from mammalian cells, APB-1 as well as CHC-1 vesicles underwent kiss-and-run on SNX-1 compartments (Fig. 1c, Suppl. Movie 1). Additional signals of both AP-1 and clathrin were observed on stationary compartments (Fig. 1c, Suppl. Movie 1), indicating that they could probably also play a role in organizing the cargo prior to being picked up by vesicles. We also noticed the formation of new clathrin vesicles from SNX-1 compartments (see Fig. 1c bottom row, Suppl. Movie 1). They formed by a mechanism of SNX-1 tubulation with subsequent recruitment of clathrin and pinching off from the tube, which retracted afterwards, reminiscent of the generation of retrograde transport carriers previously observed (Cullen and Steinberg, 2018; Gopaldass et al., 2024; Naslavsky and Caplan, 2018). Clathrin was also frequently associated with RAB-5 early endosomes (Suppl. Fig. 1b).

Activation of AP-1 requires Arf1. Moreover, we and others have provided evidence for a role of ARF-1 GTPase in endosomal recycling (Ackema et al., 2013; Ishida et al., 2023; Kawada et al., 2015; Stockhammer et al., 2024). To study the localization and dynamic behavior of ARF-1 in *C. elegans*, we established endogenously mNeongreen (ARF-1-mNG) or mScarlet-I (ARF-1-mSc) tagged ARF-1 strains using CRISPR/Cas9 technology. A fraction of ARF-1 was associated with FERARI positive domains (Fig. 1b) on SNX-1 compartments (Fig. 1g), where ARF-1 appeared to cluster in nanodomains. Interestingly, docked vesicles with CHC-1, hTfR and RAB-11 were frequently found near or at these ARF-1 nanodomains, indicating an involvement of ARF-1 in the kiss-and-run process (see Fig. 1d-f line graphs). We were wondering, whether this function is conserved in mammalian cells. Indeed, Arf1 co-localized with Rab11 and SNX1 at sites where Rab11 vesicles docked on the sorting endosomes (Fig. 1h), suggesting evolutionary conservation. Most importantly, ARF-1 was present at the FERARI-mediated KAR sites between vesicles and sorting compartments (Fig. 1d-h). Our data indicate that ARF-1 could be either present on the vesicle or on the sorting compartment, or both. To address this question, we analyzed kiss-and-run movies with either hTfR or RAB-11 as vesicle markers, SNX-1 and ARF-1 (Fig. 2). Strikingly, we detected specific ARF-1 signals that appeared only for the duration of the kiss at SNX-1 compartments for both hTfR (Fig. 2a, Suppl. Movie 2) and RAB-11 (Fig. 2b, Suppl. Movie 2) vesicles. These data suggest that ARF-1 might be activated for a short period of time during kiss-and-run at KAR sites. Taken together, these data indicate that ARF-1 is present at the KAR site between RAB-11 vesicles and SNX-1 structures, and we hypothesize that ARF-1 may serve evolutionary conserved important functions during kiss-and-run.

**Figure 2:**
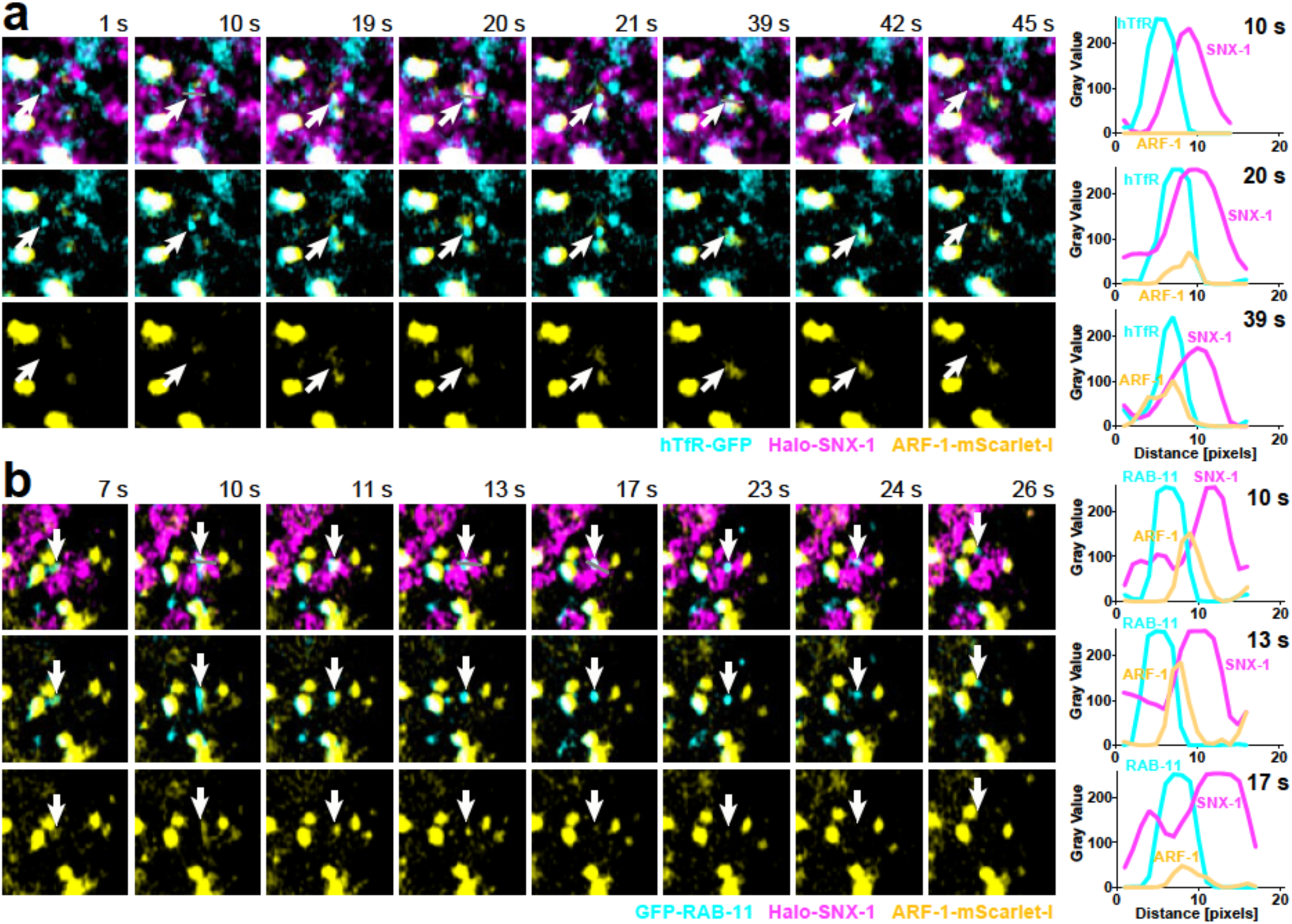
ARF-1 is present during kiss-and-run events at SNX-1 compartments. (a) Cargo vesicle with hTfR during kiss-and-run interaction with SNX-1 compartment. Arrow points at vesicle position. For clarity, SNX-1 (middle) and also hTfR (bottom) were removed to show the presence and short flare up of ARF-1 on and near the vesicle during the docking. In selected frames, the presence of ARF-1-mScarlet-I was measured with line plots shown on the right. (b) Same experiment as (a) but with RAB-11 vesicle.

### Reducing ARF-1 activity on endosomes causes defects in TEN formation

If ARF-1 plays an important role during kiss-and-run, loss of ARF-1 should affect the SNX-1 compartment. This is indeed, what we observed. Knock-down of ARF-1, or its paralog ARF-5, had a dramatic effect and caused the accumulation of short, stiff SNX-1 tubules in intestinal cells, that we called "needles" (Fig. 3a-e, Suppl. Movie 3). This needle phenotype is conserved in *S. cerevisiae*. Yeast strains harboring the inactive *arf1-18* allele displayed elongated, needle-like endosomal compartments (Fig. S1c). We had previously shown that the ArfGEF GBF-1 is required for efficient endosomal transport in oocytes and the intestine in *C. elegans* (Ackema et al., 2013). Therefore, we knocked-down GBF-1, which resulted in the same needle phenotype. RNAi of *arf-1* and *arf-5* showed a somewhat weaker effect than *gbf-1*, potentially due to some redundancy in function. These needles were still positive for FERARI (Fig. 3a and b), indicating that recruitment of the FERARI machinery to the SNX-1 positive structures is independent on ARF activity. To test whether the needles are still active in recycling, we analyzed the localization of the early endosomal RAB-5 and recycling endosomal RAB-11. Both, RAB-5 and RAB-11 structures docked onto the needles (arrows in Fig. 3c and d). However, these vesicles were less mobile and were often affixed on the needles in the absence of ARF-1, ARF-5 or GBF-1, suggesting that they could fuse with, but not leave, the needles - they could kiss but not run (Supple. Movie 3). Consistent with this notion, the cargo hTfR was detected in vesicles and in the needles under knockdown conditions (Fig. 3e). Interestingly, the hTfR cargo was more often found along the needles, while RME-1, RAB-11 and RAB-5 were mostly localized at the ends (Fig. 3f). Moreover, AP-1 and clathrin were still recruited to the SNX-1 needles (Fig. 3g). Sometimes, several clathrin dots were found on a single SNX-1 needle (see arrows in Fig. 3g). In contrast to these small clathrin vesicles there are also larger clathrin compartments, which often associate with RAB-5 and are not moving (Fig. S1b). Our data so far indicate that the tethering function of FERARI is still intact, but the fission function might be altered in the absence of GBF-1, ARF-1 or ARF-5. Since GBF-1 is the activator of ARF-1 and ARF-5, we assumed that GBF-1 should be close by and might even interact with FERARI. Indeed, in HeLa cells, GBF1 was found in close proximity to the FERARI subunit EHD1 on SNX1 positive endosomal structures (Fig. 3h). Moreover, the N-terminal domain of GBF-1 interacted with the ankyrin repeats of UNC-44 (the *C. elegans* ANK1-3 homolog) in a yeast two-hybrid assay (Fig. 3i, Suppl. Fig. 1d, UNC-44 domains are shown in Fig. 5d). In support of our findings, interactions between GBF1 and Rab11A, Rab5A, SNX1 and SNX2 have been reported previously (Oughtred et al., 2021). We hypothesize that the lack of ARF activation will lead to the formation of short endosomal intermediates in the sorting process (as seen in Fig. 1c bottom) because tubules are still formed from Rab5 positive-sorting endosomes, to which FERARI, cargo and other factors can be recruited but the recycling process is blocked and hence needles accumulate and the TEN cannot be formed.

**Figure 3:**
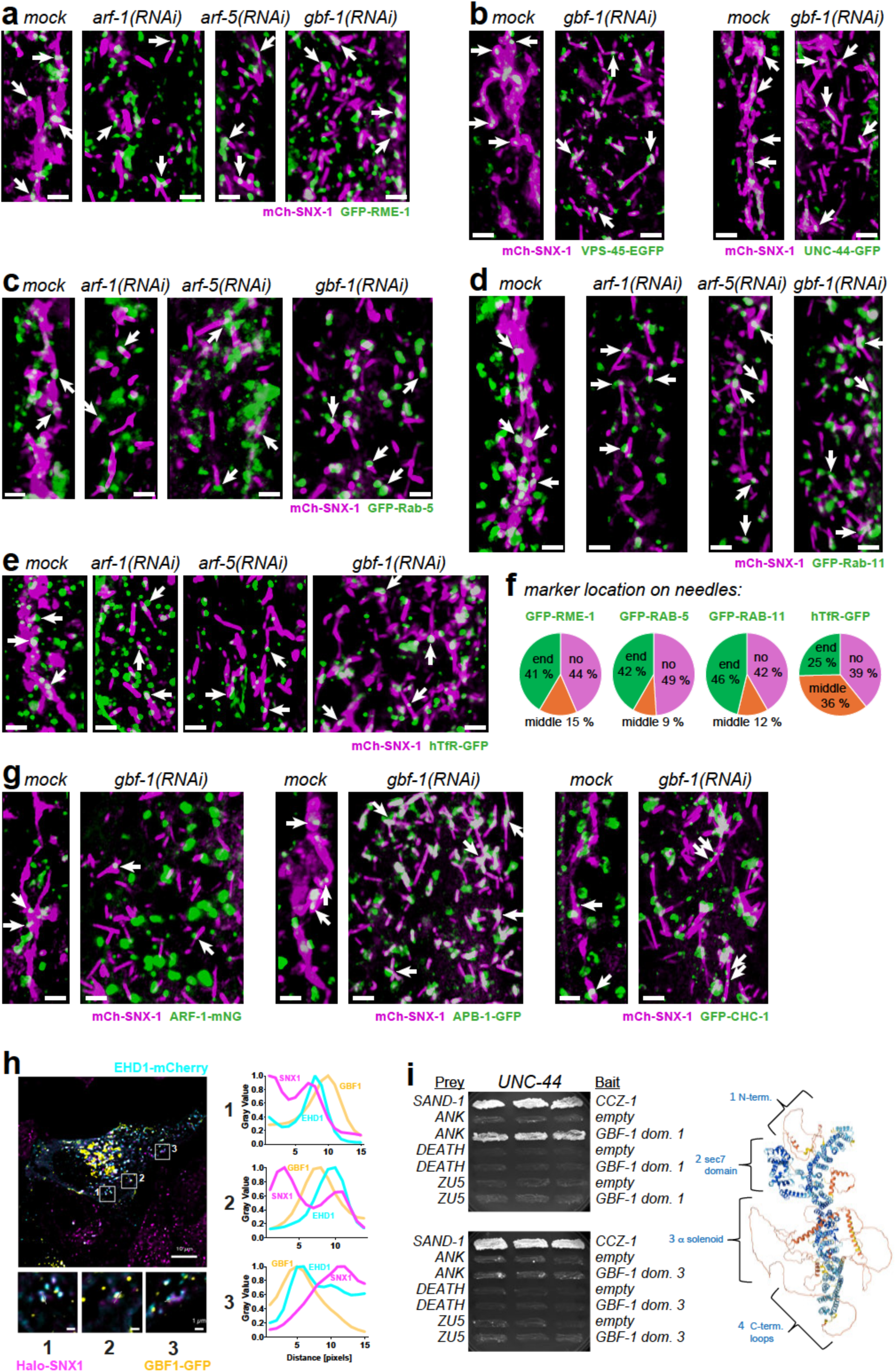
Deactivated ARF-1 causes short SNX-1 tubules that are connected to RAB-5, RAB-11 and contain cargo and RME-1. (a) Knock-downs of *arf-1*, *arf-5* and *gbf-1* lead to formation of SNX-1 short tubules or needles. Often, these short tubules are connected to RME-1 compartments (arrows). Shown are 3D projections of relevant compartments on the apical side of intestinal cells. (b) VPS-45 and UNC-44 behave in a similar way as RME-1 in *gbf-1(RNAi)*. (c) Short tubular SNX-1 compartments in *arf-1*, *arf-5* and *gbf-1* RNAi are often connected to RAB-5 vesicles. (d) RAB-11 is also closely associated to the SNX-1 short tubules caused by *gbf-1(RNAi)*. (e) Short SNX-1 needles contain hTfR cargo (arrows). Scale bars are 2 µm in all of figure 3. (f) Quantification of signal position of markers used in (a), (c), (d) and (e) on needles. (g) Localization of ARF-1, APB-1 and CHC-1 on SNX-1 compartments in *gbf-1* knock-down strains. Arrows point to domains or vesicles closely associated with SNX-1 needles. Scale bars are 2 µm. (h) GBF1 and EHD1 can be observed in close proximity to SNX1 compartments in HeLa cells. Enlargements show three examples with line plots demonstrating the overlap between GBF1 and EHD1 as well as the close association with SNX1 (graphs on the right). (i) Interaction of GBF-1 with FERARI subunit UNC-44 shown by yeast two-hybrid assays. GBF-1 was subdivided into 4 domains shown on the right. Interactions with domains 1 and 3 are shown, for domains 2 and 4 see Fig. S1d.

### AGEF-1 activates ARF-1 on TEN

We saw striking effects on sorting endosomes by knocking down *gbf-1*, indicating that GBF-1 might be required for the formation of the TEN and FERARI mediated kiss-and-run on sorting endosomes. However, in mammalian cells the AGEF-1 homologs BIG1 and BIG2 are mostly thought to act on endosomes (Boal and Stephens, 2010; Ishizaki et al., 2008; Shin et al., 2004). We investigated the role of AGEF-1 in recycling by performing a knockdown in a SNX-1::mCherry strain. We observed a dramatic enlargement of SNX-1 compartments (Fig. 4a). Again, comparable to earlier results with *gbf-1*, we observed that RAB-5 was not much affected while RAB-11, RME-1, hTfR and APB-1 accumulated to great extent on these enlarged structures (Fig. 4a). Perhaps not surprisingly, ARF-1 was absent from these structures. These data indicate that *agef-1(RNAi)* causes a traffic jam at SNX-1 compartments, on which cargo and cargo adaptors are stalled probably at KAR sites. Since the structures are grossly enlarged compared to normal, we surmise that this block might happen downstream of GBF-1 mediated TEN formation. The TEN would be filled with cargo that cannot flow into RAB-11 vesicles fused at KAR sites. We also observed direct interactions between AGEF-1 domains with UNC-44 and RME-1 (Fig. 4b and c, Suppl. Fig. 1e, f). Interestingly, BIG1 and BIG2 were found to interact with Rab11A (Oughtred et al., 2021). Our data indicate that ARF-1 might play a dual role in recycling to the plasma membrane. ARF-1 would be required first for the formation of TEN and then play a role in cargo loading into RAB-11 recycling vesicles. We surmise that the activation of ARF-1 on sorting endosomes is dependent on GBF-1, while the activation of ARF-1 on TEN requires AGEF-1 activity (Fig. 4d). We decided to concentrate on the role of GBF-1 in FERARI-dependent recycling.

**Figure 4:**
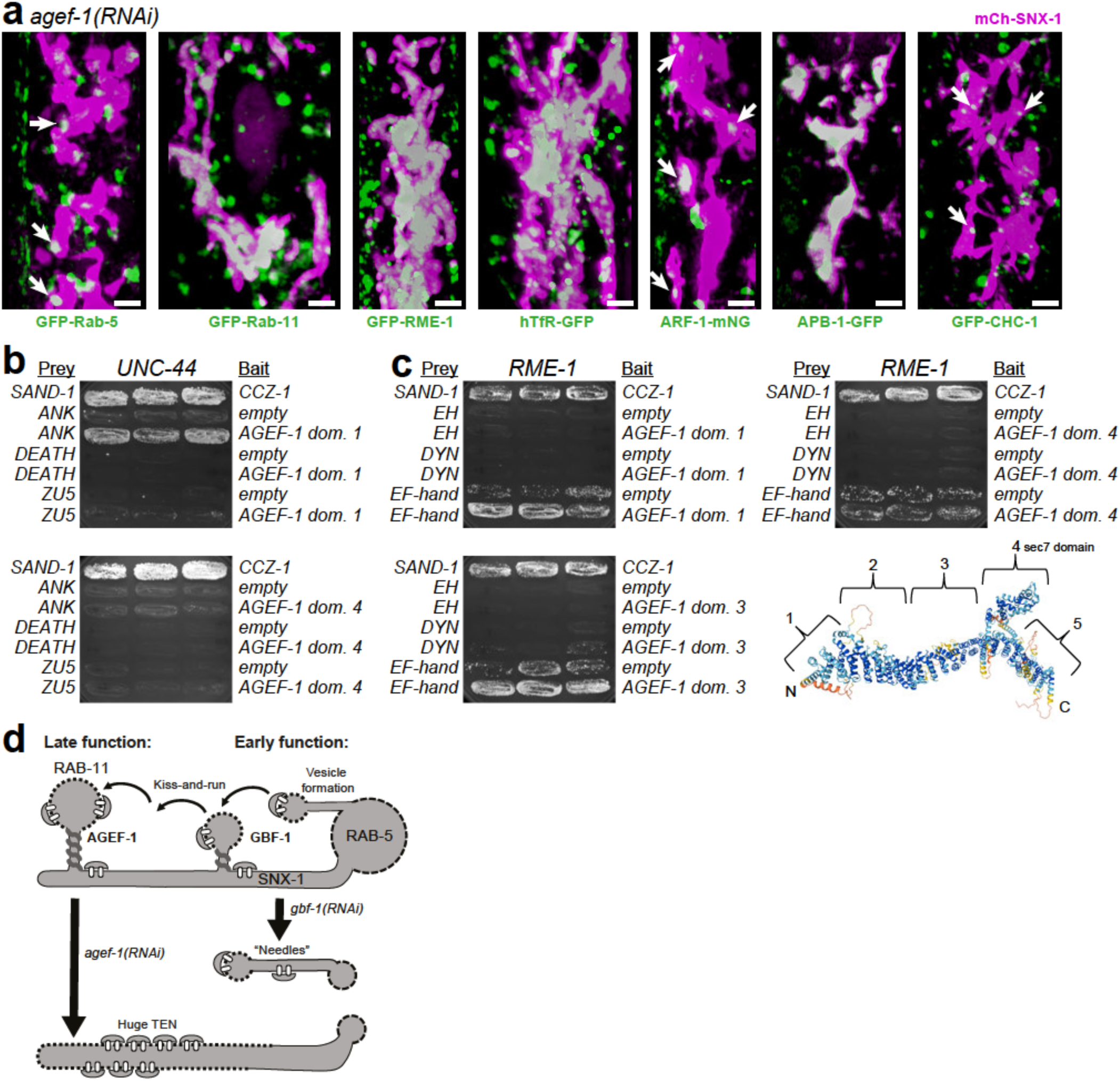
Loss of AGEF-1 causes a major block in exit from TEN with giant SNX-1 compartments. (a) 3D projections of enlarged SNX-1 compartments showing some close association of RAB-5, ARF-1 and CHC-1 (clathrin) (indicated by arrows). The other markers RAB-11, RME-1, hTfR and APB-1 accumulate in large bright patches. Scale bars in figure are 2 µm. (b) Interaction of AGEF-1 domains with UNC-44 in yeast two-hybrid experiments. Domain 1 shows interaction, while the sec7 domain does not. The other domains showed very weak growth in the assay (see Fig. S1e, f). (c) Interaction of AGEF-1 with EF-hand domain of RME-1. Domains 1 and 3 showed robust interaction, while the GEF domain did not interact (the other domains were only partially growing as shown in Fig. S1e, f). Schematic representation of AGEF-1 domains is shown on the right. (d) Hypothetical functions of AGEF-1 and GBF-1 during endosomal recycling. Late function of AGEF-1 may lead to large SNX-1 intermediates, while an early involvement of GBF-1 results in short needle-like TEN with RAB-5 and RAB-11 at ends.

**Figure 5:**
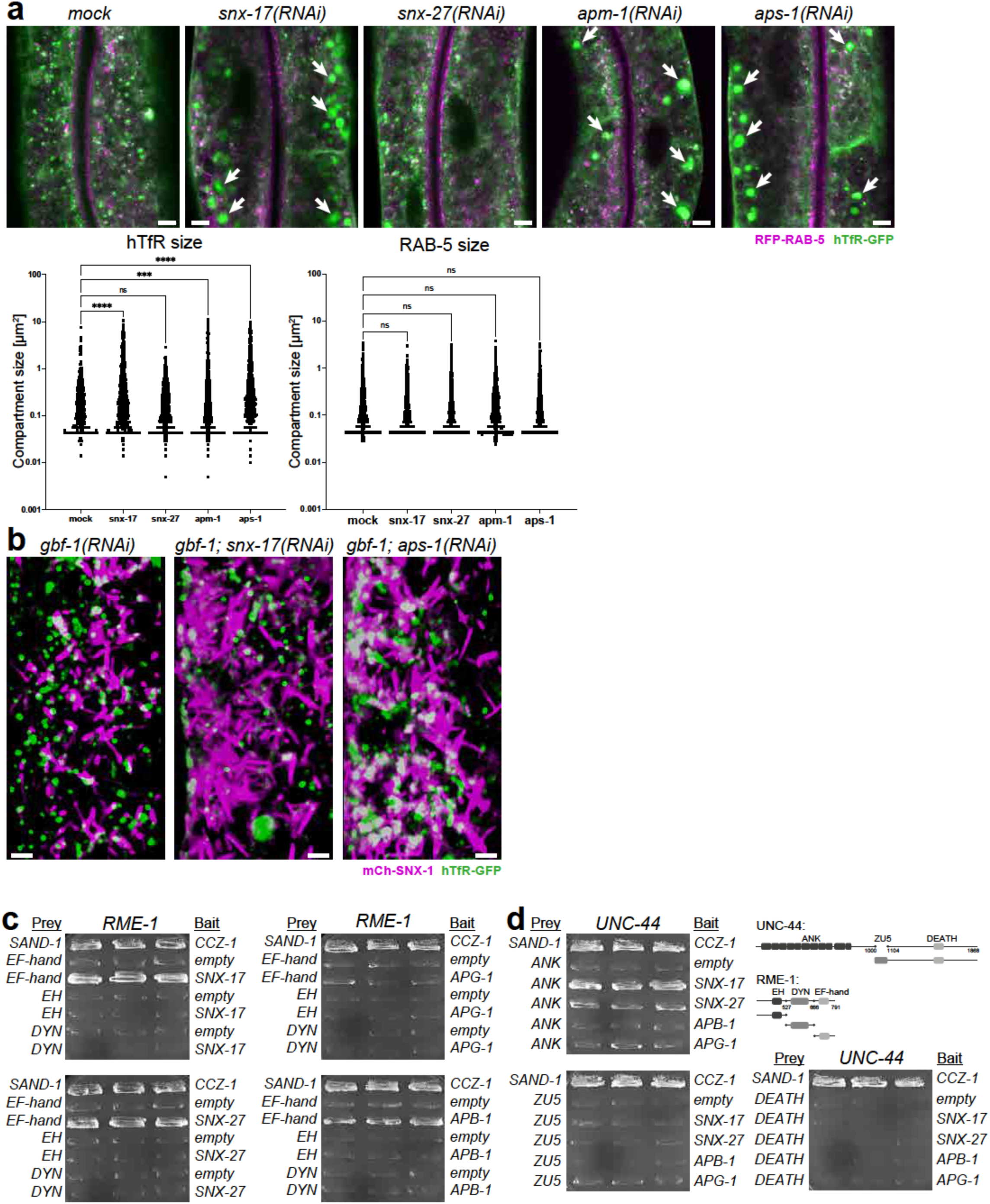
Traffic jam of cargo and cargo adaptors in SNX-1 disrupted tubules formed by *gbf-1(RNAi)*. (a) Knock-down of cargo adaptors *snx-17, apm-1* and *aps-1* cause accumulation of hTfR, but do not affect RAB-5 positive sorting endosomes. Loss of *snx-27* has no effect on hTfR recycling. 5 µm scale bar. Quantifications of compartment sizes are shown below. (b) Cargo adaptors *snx-17* and *aps-1* have differential effects on hTfR sorting into SNX-1 needles in a *gbf-1(RNAi)* background. Scale bars are 2 µm. (c) Cargo adaptors interact with FERARI. Yeast two-hybrid analyses show interactions between SNX-17, SNX-27 and APB-1 with the EF-hand motif of RME-1. (d) SNX-17 and SNX-27 also bind to the ankyrin repeats of UNC-44. The domains used in the yeast two-hybrid experiments are shown on the right.

### The role of cargo adaptor activation by ARF GTPases during endosomal sorting

We next aimed to elucidate the role of ARF-1 in cargo loading at SNX-1 sorting compartments. We have previously shown that cargo adaptors and cargo availability play an important role in determining the lengths of the kiss of RAB-11 endosomes at KAR sites (Solinger et al., 2020; Solinger et al., 2022). In particular, the sorting nexins SNX-17 and SNX-27, and the adaptor complex AP-1 seemed to be crucial for effective recycling. Arf1 has been shown to activate AP-1 in mammals (Ren et al., 2013) providing a possible link between cargo loading and ARF-1 activity. We performed knock-downs of *snx-17, snx-27* and the AP-1 complex subunits *apm-1* and *aps-1* and assessed their effect on the cargo hTfR and RAB-5 (Fig. 5a). Similar to what we had observed previously, hTfR was blocked in enlarged SNX-1 positive compartments in *snx-17* and AP-1 knock-downs but was not affected by *snx-27(RNAi)* (Solinger et al., 2022). In contrast, the RAB-5 compartment was not perturbed by the transport block, indicating that the cargo accumulated in compartments distinct from early endosomes (Fig. 5a). Next, we tested our hypothesis that ARF-1 function is required for cargo flow in the recycling pathway. To this end, we performed double knock-downs of *gbf-1* together with either *snx-17* or *aps-1* (Fig. 5b). The *gbf-1(RNAi)* needle phenotype was greatly exacerbated in the double knockdowns, supporting a potential role of active ARF-1 in recycling from sorting endosomes to the plasma membrane. The hTfR accumulation was less pronounced in *gbf-1; snx-17(RNAi)* animals compared to the *gbf-1; aps-1(RNAi)* animals. This observation is consistent with the notion that SNX17 protects cargoes from lysosomal degradation (McNally et al., 2017; Steinberg et al., 2012), while AP-1 plays a major role in recycling cargo from endosomes to the plasma membrane and the Golgi (Buser and Spang, 2023; Robinson et al., 2024; Solinger et al., 2022; Uemura et al., 2021). If ARF-1 was indeed required for the process, we would assume that in *gbf-1(RNAi)*, ARF-1 would be lost from the SNX-1 compartment and the Golgi, GBF-1’s canonical site of function. As expected, ARF-1 no longer co-localized with SNX-1 or the Golgi marker Mans (Fig. 3g left and Fig. S2a, b). We hypothesized that in the absence of active ARF-1 cargo flow from the needles into the recycling vesicles is dysregulated and hence fission of the stalk between the vesicle and the needle cannot occur. If this was the case, one would expect a link between cargo adaptors and FERARI, which would be responsible for membrane fission. We performed yeast two-hybrid experiments to detect direct interactions between cargo adaptors and FERARI. SNX-17, SNX-27 and the ß-subunit of AP-1, APB-1, but not AP-1’s γ subunit APG-1, interacted with the pinchase subunit of FERARI, RME-1 (Fig. 5c). The binding site resided in all cases in the C-terminal domain of RME-1, which contains an EF-hand, an established protein-protein interaction domain (Lewit-Bentley and Rety, 2000).

Similar interactions between the mammalian counterparts SNX17 and EHD1 have been described (Dhawan et al., 2020). In addition, we also observed interactions of SNX-17 and to a lesser extent SNX-27 with the ankyrin repeats of UNC-44 (Fig. 5d). The different interactions of SNX-17 and SNX-27 on one side and AP-1 on the other might correlate with the differential requirements of SNX-17 and AP-1 in the recycling process. The direct interaction between cargo adaptors and FERARI at the KAR sites is consistent with our model of cargo flow between recycling carriers and the SNX-1 sorting compartment.

### The cargo adaptor SNX-6 acts as a cargo coordinator in the SNX-1 sorting compartments

These cargo adaptors are all found mostly on transport vesicles and are recruited to vesicle exit or KAR sites. In contrast, SNX-6 is a cargo adaptor that is also largely found on the tubular part of sorting endosomes and TEN (Lopez-Robles et al., 2023). We have previously shown that several types of vesicles (RAB-5, RAB-11 and RAB-10) were blocked on SNX-1 tubules upon *snx-6* knock-down (Solinger et al., 2022). This is in a way reminiscent of the *gbf-1(RNAi)* phenotype, where vesicles were also blocked on the TEN. Since SNX-6 plays an important role in organizing the cargo for proper flow into vesicles, we determined the effect of *snx-6* depletion on cargo flow through SNX-1 networks. Knock-down of *snx-6* leads to accumulations of hTfR in SNX-1 networks (Fig. 6a). Also, SNX-1 networks are smaller and less prevalent, since SNX-6 forms a heterodimer with SNX-1 and provides structural integrity to sorting tubules (Simonetti et al., 2019). We previously observed that vesicles are blocked under these conditions and docking is prolonged during kiss-and-run (Solinger et al., 2022). Moreover, as expected, hTfR accumulated in the SNX-1 compartment. We obtained similar results with RAB-11, that also accumulated upon *snx-6(RNAi)* (Fig. S2c). ARF-1 appeared to be more often associated with SNX-1 in cells lacking *snx-6* compared to control (Fig. 6b). Similar accumulations of clathrin (CHC-1) on SNX-1 tubules in *snx-6(RNAi)* were observed, suggesting that both might be present at the same site on the SNX-1 tubule. Moreover, since the flow of cargo is inhibited, APB-1 accumulated and covered the entire SNX-1 network (Fig. 6b). Note that the pattern of hTfR and APB-1 was not the same (compare Fig. 6a to 6b), indicating that hTfR is segregated into distinct domains within the SNX-1 tubules.

**Figure 6:**
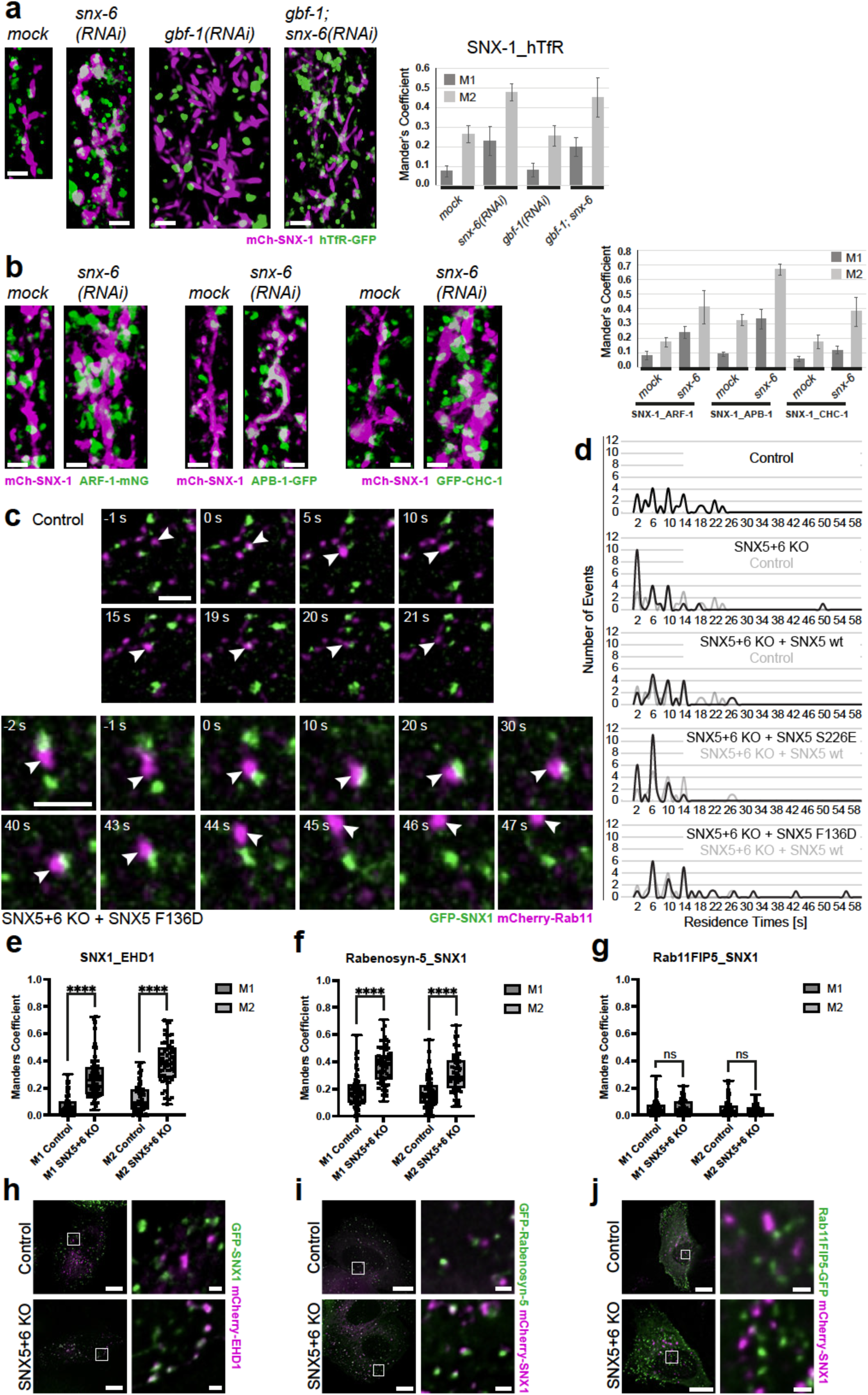
Cargo coordination by **SNX-6 cargo adaptor (ESCPE) facilitates cargo flow into RAB-11 vesicles.** (a) Cargo hTfR accumulates in SNX-1 sorting tubules upon *snx-6(RNAi)*. It also accumulates on *gbf-1(RNAi)* induced SNX-1 needles upon *snx-6* knock-down. Mander’s coefficients are shown on the right, with average and standard deviations (n=5). (b) Increased ARF-1 and CHC-1 association with SNX-1 tubules in *gbf-1* knock-down intestinal cells, while APB-1 almost completely covers SNX-1 sorting compartments. Co-localization was measured and is shown on the right (average and standard deviations, n=5). Scale bars are 2 µm. (c) Examples of Rab11 kiss-and-run movie stills (Movie 4) for control and non cargo binding SNX5 mutant. Scale bars are 2 µm. (d) Residence times of kiss-and-run with Rab11 vesicles in HeLa cells expressing SNX5 mutant variants as indicated. (e) SNX5+6 double knock-out cells show increased co-localization of SNX1 and EHD1. (f) SNX5+6 double knock-out cells show increased co-localization of SNX1 and Rabenosyn-5. (g) No increase in co-localization of SNX1 and Rab11FIP5 upon SNX5+6 knock-out. (h) Representative cells and enlargements for (e). (i) Representative cells and enlargements for (f). (j) Representative cells and enlargements for (g). Scale bars are 10 µm for cell overview and 1 µm for inlays.

To corroborate and extend the role of SNX-6 in cargo flow, we turned to mammalian cells. In mammalian cells, SNX5 and SNX6 fulfill the function of *C. elegans* SNX-6. We used a cell-line in which both SNX5 and SNX6 were deleted (Solinger et al., 2022) and in which we expressed either wild-type SNX5, a mutant that could not bind to sorting endosomes (SNX5 (S226E)) or a mutant defective in cargo binding (SNX5 (F136D)) (Simonetti et al., 2019). We then determined the residence time of Rab11 (Fig. 6c, d, Suppl. Fig. 3a, b, Suppl. Movies 4, 5). As expected, SNX5/SNX6 double knock-out led to a reduction in the residence time of Rab11 vesicles at the SNX1 compartments, which was rescued by re-expression of wild-type SNX5. In contrast, the SNX5 (S226E) mutant, which cannot be efficiently recruited to the sorting compartment, did not rescue the reduced residence time phenotype. However, the SNX5(F136D) mutant that abolishes cargo binding caused even longer docking times of vesicles (Fig. 6c, d, Suppl. Fig. 3a, b). Additionally, SNX5+6 knock-out caused an increase in co-localization between FERARI subunits EHD1 and Rabenosyn-5 with SNX1 compartments, but not for Rab11FIP5 (Fig.6 e-j). This is in agreement with previous results showing that vesicles are blocked during kiss-and-run when SNX5 and SNX6 are missing. Probably, Rab11FIP5 is primarily associated with the incoming Rab11 vesicles (Suppl. Fig. 3c). These experiments underscore the important role of SNX5/SNX6 (also known as ESCPE) for kiss-and-run and cargo sorting in the SNX-1 sorting compartment and their evolutionary conservation from worms to mammals. In summary, the function of SNX-6 appears to be in coordinating the cargo flow probably retaining cargo in SNX-1 sorting compartments and preventing back-flow into the fused vesicle.

### ARF-1 over-activation causes SNX-1 sorting compartments to be blocked as globules

Our data thus far indicate that activation of ARF is crucial for proper cargo loading into vesicles. Therefore, we asked next whether over-activation of ARF-1 might accelerate the recycling process. To this end we knocked-down the *C. elegans* ArfGAP1 and ArfGAP2 homologues RFGP-1 and RFGP-2, respectively. To our surprise, the flow through the SNX-1 compartment was still blocked in *rfgp-1* and *rfgp-2* knock-downs. SNX-1 tubular networks disassembled into globular structures (Fig. 7a). To ascertain that ARF-1 was over-active, we carried out *rfgp-1* and *rfgp-2* knock-downs in the ARF-1-mNeonGreen strain. Under these conditions, the SNX-1 structures were indeed marked by ARF-1 (Fig. 7c, Fig. S2b). These globules were sometimes still connected to RAB-5 compartments but were not RAB-5 positive overall (Fig. 7a). On the other hand, the SNX-1 globules were very often RAB-11- and AP-1 positive (Fig. 7a, c, d, Fig. S3d). In contrast, clathrin was not enriched on these structures (Fig. 7c). Similar effects were observed in yeast strains harboring either the hyperactive *arf1-11* or the hypoactive *arf1-18* allele at the permissive temperature (Fig. S1c). We observed that *arf1-11* cells exhibited enlarged endosomal compartments while largely maintaining a round morphology, comparable to the control strain. In contrast, *arf1-18* cells displayed significantly elongated, needle-like endosomal compartments while retaining an area comparable to the control. Thus, ARF-1 over-activation rather seemed to block recycling at a stage in which RAB-11, AP-1 and SNX-1 co-localize at sorting compartments and potentially also interfere with TEN formation. As in the case of *gbf-1(RNAi)*, FERARI was present on the SNX-1 compartment, which contained also the cargo hTfR (Fig. 7b). Therefore, ARF-1 activity is not essential for FERARI recruitment and cargo deposition into the SNX-1 compartment. FERARI might rather coordinate ARF-1 activity because the FERARI subunit UNC-44 interacted directly with both the ArfGEFs GBF-1 and AGEF-1 and the ArfGAP RFGP-2 (Fig. 3i, 4b and 7e). In mammalian cells, interactions of ArfGAP2 with VPS45 haven been reported previously (Oughtred et al., 2021). Moreover, RFGP-2 also bound to the FERARI pinchase subunit RME-1 (Fig. 7e). The binding sites of GBF-1 and RFGP-2 might overlap preventing simultaneous binding to UNC-44. Our data are consistent with the requirement of ARF-1 cycling between the inactive and active state for efficient cargo sorting into RAB-11 recycling vesicles destined to the plasma membrane.

**Figure 7:**
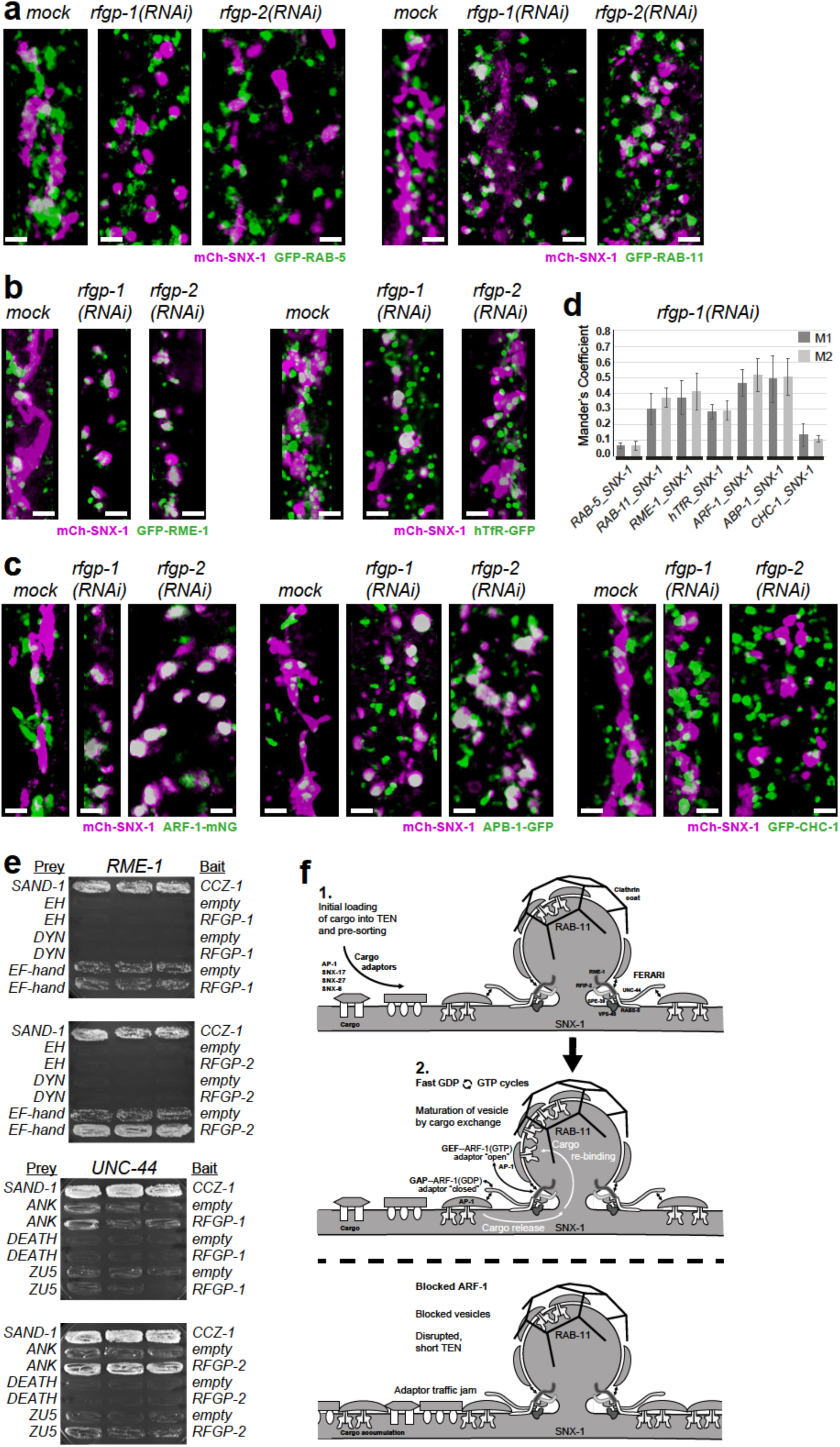
Over-activated ARF-1 causes engorged short SNX-1 globules with accumulations of RAB-11, RME-1, cargo, ARF-1 and APB-1, but not RAB-5 and clathrin. (a) Knock-downs of *rfgp-1* or *rfgp-2* lead to disruption of SNX-1 tubules and formation of globular TEN that contain RAB-11 (right), but are only rarely connected to RAB-5 (left). (b) Engorged SNX-1 globules in *rfgp-1* or *rfgp-2(RNAi)* strains recruit RME-1 (FERARI subunit) (left) and contain hTfR cargo (right). (c) ARF-1 accumulates on SNX-1 globules in the absence of *rfgp-1* or *rfgp-2* (left). While this leads to an over-recruitment of APB-1 (probably AP-1) (middle) it does not block clathrin (CHC-1) on these structures (right). Scale bars throughout figure are 2 µm. (d) Mander’s coefficients measured for (a)-(c) (n=5 worms each), average and standard deviations are shown. (e) ArfGAP interactions with FERARI. Yeast two-hybrid analyses show interactions between RFGP-2 and the EF-hand motif of RME-1 and the ankyrin repeats of UNC-44 (domains are shown in Fig. 5d). No interaction was seen with RFGP-1 and these FERARI subunits. (f) Hypothesis for cargo loading into vesicles at sorting endosomal SNX-1 tubules (see also Movie 6). 1. Under normal conditions, cargo adaptors AP-1, SNX-17 and SNX-27 will corral cargo with similar transport destinations and through binding to FERARI, bring the cargo into close proximity to vesicle docking sites (KAR sites on the TEN). 2. Because FERARI forms a diffusion barrier for cargo flow into vesicles, the cargo adaptors on SNX-1 need to dissociate from the cargo and hand it over to cargo adaptors on the vesicles, thereby passing through the structure needed for vesicle docking and stabilization. This cargo shuffling can be achieved by fast GDP-to-GTP cycles of ARF-1 that regulate the binding of cargo to AP-1. Cargo adaptors on the vesicle will be activated to bind cargo and to release FERARI when cargo is bound, thereby facilitating the pinching-off of the vesicle. Bottom: Without actively cycling ARF-1 (either by de-activation with *gbf-1(RNAi)* or by constant over-activation through *rfgp-1/2* knock-down) cargo is blocked in SNX-1 structures. With ArfGAPs present in the *gbf-1* knock-down, the cargo adaptors will release the cargo and result in thin SNX-1 needles. Vesicle formation will be blocked, because no active cargo adaptors will bind the cargo. In the presence of GBF-1 but lacking ArfGAPs in the *rfgp-1/2* knock-downs, over-active ARF-1 will lead to accumulation of AP-1 on the SNX-1 compartments and these will be engorged with cargo. Vesicle formation will still not function, because cargo would not be able to pass through the diffusion barrier around the docking site.

## Discussion

We have previously shown that cargo flows from SNX-1 endosomes and TEN into RAB-11 recycling vesicles during kiss-and-run in a FERARI-dependent manner (Solinger et al., 2020; Solinger et al., 2022). Moreover, the amount of cargo that is available for transport to the plasma membrane and cargo receptors are critical determinants of the time the recycling vesicle is fused to the TEN. However, how this process is regulated and coordinated at KAR sites remained elusive. Here, we addressed this gap of knowledge. We hypothesized that intriguing candidates for this regulation would be ARF GTPases that can be activated by GEFs and switched off by GAPs. They have been implicated in similar cargo transfer mechanisms at the trans Golgi network (TGN) and in Golgi to ER transport (D’Souza-Schorey and Chavrier, 2006; Pankiv et al., 2024). Also, the involvement of Arf1 in endosomal transport was observed in yeast (Estrada et al., 2015; Kawada et al., 2015) where the ArfGAP Glo3 (the orthologue of RFGP-2 in *C. elegans*) is important for diverse intracellular traffic pathways. In worms, the ArfGEF GBF-1 was shown to play a role in endosomal transport (Ackema et al., 2013). In humans, Arf1 is required for the integrity of the recycling pathway and for lysosomal function (Kondo et al., 2012; Szentgyorgyi et al., 2024). Moreover, a mutation in ARF1 derived from a neurological disorder alters the morphology of recycling endosomes (Ishida et al., 2023). Our results provide strong evidence that *C. elegans* ARF-1-and ARF-5- are indeed involved in cargo loading into vesicles during kiss-and-run. Both de-activation or over-activation of ARF resulted in loss of transport through TEN, resulting in a traffic jam with cargo, cargo adaptors, FERARI machinery and Rab GTPases being recruited, but unable to continue through the pathway. Vesicles bearing clathrin were mostly unaffected by these accumulations, indicating a block before loading of cargo into these transport carriers. We propose that ARF-1 (and ARF-5) need to cycle between active (GTP bound) and inactive (GDP bound) states, for successful cargo loading (Fig. 7f, Movie 6). A similar mechanism has been proposed for cargo loading into COPI vesicles to increase cargo density in vesicles (Lanoix et al., 2001; Weiss and Nilsson, 2003). Why would such a mechanism be particularly necessary to load cargo into vesicles at KAR sites? The problem might be at least in part a question of the architecture of the machinery needed for kiss-and-run. To avoid that a vesicle fuses completely into the receiving membrane and flattens out, a protein scaffold around the fusion pore is needed, a sort of restricting collar preventing fusion pore expansion. In addition, the vesicle itself might lose structural integrity, without a coat that keeps it from deforming. A docking vesicle will therefore be stabilized by a complex protein structure consisting of FERARI (maybe with an RME-1 spiral), clathrin, probably the actin cytoskeleton and additional factors. However, this dense thicket of proteins will pose a problem for cargo transport across collar barrier and into the vesicle. This arrangement would pose a steric hindrance problem for transfer of cargo bound to cargo adaptors into the vesicle. Inclusion of cargo-cargo adaptor subcomplexes is the basis of the general model of transport vesicle generation and budding. Our data indicate that cargo-adaptor interaction already take place on the TEN, and are not restricted to cargo in the vesicle. Therefore, cargo needs to be released from the cargo adaptor on the SNX-1 compartment and transferred into the vesicle more or less one behind the other, sequentially and in a coordinated manner. We propose that breaking and reinstating cargo-cargo adaptor interaction can be achieved by ARF-1. Cycles between ARF-1-GTP and ARF-1-GDP would act as a switch to turn cargo adaptors "off" (closed, releasing cargo) on the TEN side and "on" (open, re-binding cargo) on the vesicle side (Fig. 7f, Movie 6). We observed different phenotypes upon knockdown of the GEF and the GAP. This could be explained by AP-1 holding on to cargo in the SNX-1 globules due to ARF-1 over-activation, being unable to transfer the cargo into vesicles and so swelling the tubules to globular shapes when ArfGAP activity is absent. On the other hand, the SNX-1 needles would tend to release cargo because of ARF-1 deactivation in the *gbf-1* knock-down. Since the cargo could not be properly bound on vesicles in the absence of "open" cargo adaptors, it would probably flow out of the tubules and the needles would remain "slim". This Arf1-dependent switching between locked and unlocked AP-1 complexes has been proposed to be important for the regulation of traffic (Ren et al., 2013). The cargo adaptors will already be organized near the exit site, because of their interaction with cargo and FERARI. Moreover, we provide evidence that ArfGAPs for cargo release and ArfGEFs for cargo binding can interact with FERARI. The concentration of cargo, their adaptors, ARF-1 and its regulators and FERARI at KAR sites would allow them to perform their respective roles in a coordinated way.

Our data reveal at least two distinct functions for ARF-1 in endosomal recycling, one sorting endosomes and one on the TEN. The function on sorting endosomes requires ARF-1 activation by GBF-1 and on the TEN by AGEF-1. Our data indicate that GBF-1 is essential for TEN formation by promoting the formation of RAB-11 recycling carriers from RAB-5 positive sorting endosomes; a process, which has also been observed in mammalian cells (Cullen and Steinberg, 2018; van Weering et al., 2012). In the absence of GBF-1, the tubes from which the RAB-11 recycling carrier bud off are still generated and also contain cargo, but they seem to be defective in that they cannot merge into a TEN and a hence give rise to the needle phenotype. It is conceivable that the needles contain somewhat less cargo or that biophysical properties of the needles are different as they seem to be very stiff and clearly do not bend or round up (Movie 3). The second function of GBF-1 would be its involvement in efficient cargo loading into recycling vesicles on sorting endosomes. We surmise that this latter function is taken over by AGEF-1 on the TEN because in the absence of AGEF-1 the TEN loses it structure and rounds up. We assume that this is due to the strong reduction of cargo efflux from the TEN, while cargo influx would be unperturbed.

It is interesting to note that GBF-1 and ArfGAP1/2 are both required for proper recycling vesicle maturation. This is reminiscent of the Golgi to ER transport module where ArfGEF and ArfGAP are sequentially needed in the process of vesicle formation and trafficking (D’Souza-Schorey and Chavrier, 2006). Perhaps the GEFs play also a role in the recruitment of the appropriate GAPs indicating that there might be more crosstalk between GEFs and GAPs than previously anticipated.

We envision a serial binding scheme for the cargo transfer into vesicles at KAR sites: 1. Cargo adaptors and cargo are recruited to FERARI exit sites. 2. ArfGAP binding will de-activate ARF-1 and cause a release of cargo from the cargo adaptor. The cargo diffuse into the vesicle. 3. On the vesicle side, an ArfGEF will activate ARF-1 which promotes cargo capture by adaptors. 4. When no cargo awaits transport or the recycling vesicle has reached its capacity and no cargo is stuck in the stalk between vesicle and the TEN, FERARI promote the pinching of the stalk and the vesicle is released. The maximal capacity of cargo in the vesicle could be determined by the geometry of the clathrin coat. Still some open questions remain. In particular, the role of the cytoskeleton in both tube formation and in kiss-and-run has not been explored yet. Therefore, more studies are needed to elucidate the coordination and the biophysical implication of FERARI-mediated kiss-and-run of RAB-11 recycling carriers on sorting endosome and the TEN.

## Methods

### Worm husbandry

*C. elegans* worms were grown and crossed according to standard methods (Brenner, 1974). RNAi was carried out as previously described (Solinger et al., 2014). All experiments were done at 20°C, and worms were imaged at the young adult stage (with only few eggs). To achieve adult worms for these analyses, *gbf-1 and agef-1* RNAi bacteria had to be diluted so that proper development could occur and this leads to a partial knock-down and some ARF-1 activation in the observed gut cells. Overall, the data is consistent with a strong reduction of ARF-1 activation.

The following worm strains and transgenes were used in this study: *arf-1::mNeonGreen (syb8159) III, arf-1::mScarlet-I (syb8122) III, Halo::snx-1(syb6782) X pwIs782[Pvha-6::mCherry::SNX-1], pwIs621[vha-6::mCherry-RME-1] X, pwIs72[vha6p::GFP::rab-5 + unc-119(+)], pwIs90[Pvha-6::hTfR-GFP; Cbr-unc-119(+)], pwIs69[vha6p::GFP::rab-11 + unc-119(+)], pwIs87[Pvha-6::GFP::rme-1; Cbr-unc-119(+)], pwIs846[Pvha-6-RFP-rab-5; Cb unc-119(+)], oxSi381[Pdpy-30::apb-1(cDNA)::GFP; unc-119(+)] II, dkIs8[vha-6p::GFP::CHC-1], tmEx144 [vps-45(+)::EGFP, rol-6d], unc-44(ju1413[unc-44::gfp::LoxP::3xFLAG]) IV*.

All “Is” markers denote stably integrated arrays, the exception is *tmEx144*, which was not integrated but showed a very high transmission and expression throughout the intestine. All promoters showed good expression in gut cells.

### Microscopy of *C. elegans*

Live microscopy on worms was performed as described (Solinger et al., 2014; Solinger et al., 2022). In short, worms were immobilized on 2% agarose pads on microscopy slides using levamisole (50 mM); cover slips were sealed using Vaseline. High resolution 3D images and movies were obtained with a Zeiss LSM 880 microscope with Airyscan capabilities. The fast mode in the Zen Black software was used for all images. For movies, resolution was traded for speed by reducing the averaging to 2-4x, resulting in the required frame speeds of 0.5-1.0 seconds to follow vesicles. To catch high enough numbers of vesicles, a region of approximately 70 µm^2^ was covered (about 2 intestinal cells). Movement could be observed up to 30-45 min after immobilization of worms. From these overview movies, smaller regions of 70 x 70 pixels were selected, showing only 1-2 vesicles and events. Since vesicles and tubules are very thin, they often appear only slightly above background. Brightness and contrast were adjusted to allow visibility of the faint vesicles and to avoid random background noise that will interfere with the visualization of the processes. This may cause the images to appear oversaturated, but the original movies were taken at low laser settings to avoid bleaching and so oversaturation was never a problem. These movies were then quantified. We observed persistent movement in many cells, and events were mainly limited by the use of only one imaging plane (due to speed limitation of the microscope). The “StackReg” plugin in Fiji was used to get rid of worm shaking and drifting motions. “Bleach correction” was used to avoid irritating blinking during repeated viewing of movies (overall, bleaching was minimal due to low laser settings). Mander’s coefficients were determined using the JACoP plugin in Fiji. Images as shown in the figures consisting of z-stacks with the relevant SNX-1 compartments were used for the co-localization measurements in 3D stacks.

### Halo Ligand staining in worms

For Halo-SNX-1 staining, worms were collected the day before imaging, by washing them from the plate (containing the appropriate mock or RNAi bacteria). Worms were added to a pellet of the same bacteria and resuspended so that they would be captured in the pellet by centrifugation. Bacteria and worms were resuspended in 50 µl of S-basal medium and transferred to a 96-well plate, where 0.75 µl of 150 mM IPTG and 0.3 µl of 3 mM JF635-HaloTag (in DMSO) were added. Plate was incubated over night at 20°C in a shaking incubator inside a humid chamber. 0.5-1 hour before imaging, worms were transferred to agar plates with OP50 to remove unbound dye from gut lumen.

### Generation of CRISPR-Cas9 knock-in (KI) HeLa cells

SNX1 was targeted at the N-terminus using the guide RNA (5’-TGGAAGAAGATGGCGTCGGG<u>TGG</u>-3’, PAM sequence underlined) cloned into the SpCas9 pX330 vector (addgene plasmid #42230). For homology-directed repair (HDR) a donor plasmid was generated consisting of ∼700 bp homology arms based on the region around the start codon of SNX1, a glycine–serine linker and loxP-G418-loxP-3xALFA-Halo for insertion. For more details regarding the generation of the plasmids see (Stockhammer et al., 2024).

Wild type HeLa cells CCL-2 (cat. no. 93021013 ECACC General Collection) were transiently transfected with the gRNA and HDR plasmid using FuGENE HD (Promega) following to the supplier’s protocol. Three days after transfection, antibiotic selection with 1.5 mg/mL G418 was started and completed after approximately a week, exchanging the medium every 2-3 days. Afterwards to excise the loxP-flanked G418 antibiotic resistance cassette, cells were transfected with a Cre-recombinase (addgene plasmid #11923) using FuGENE. To further increase the KI efficiency, cells were fluorescence-activated cell sorting (FACS) using JFX_650_-CA as a Halo substrate.

### Cell culture and transient transfection

HeLa cells were cultured at 37°C and 5% CO_2_ in growth media (DMEM high-glucose medium (Sigma-Aldrich, D57969) with 10% FCS (Biowest, S1810), 1% penicillin–streptomycin (Sigma-Aldrich, P4333), 1 mM sodium pyruvate (Sigma-Aldrich, S8636) and 2 mM L-Glutamine (Gibco, 25030-024)). Cells were regularly tested for mycoplasma using a Mycoplasma PCR detection kit (Applied Biological Materials, G238).

HeLa Cells were plated before transfection at 60–70% confluency in a 6-well and transfected 6 hours later using the Helix-in transfection reagent (OZ biosciences, HX11000) according to the manufacturer’s instructions. Generally, 1 µg of DNA/plasmid was used for transfection. For the mCherry-EHD1 and GBF1-eGFP double transfection 1.95 and 0.05 µg of DNA was used, respectively. One day post-transfection cells were reseeded into a 4-well Imaging plate (Ibidi, 250228/4). Two days post-transfection cells were imaged in imaging media (DMEM high-glucose medium without phenol red (Sigma-Aldrich, D1145) with 10% FCS, 1% penicillin–streptomycin, 1 mM sodium pyruvate and 2 mM L-Glutamine). For Colocalization studies of SNX1 with FERARI subunits cells were directly seeded in a 4-well imaging plate at 60-70% confluency, transfected 6 hours later (0.26 µg of DNA/plasmid) and imaged the following day.

### Microscopy and Image analysis of HeLa Cells

Prior to imaging Halo-SNX1 KI cells (transfected with EHD1/GBF1 or Rab11/Arf1 for colocalization analysis) were stained in 200 nM JFX650 (Janelia, HT107A) in growth media for 1h at 37°C, 5% CO_2_ and subsequently washed 3 times in growth media. Subsequently, cells were left to recover in growth media for 1 hour at 37°C, 5% CO_2_. After Recovery cells were washed twice in imaging media and imaged immediately. Z-stacks of the cells were recorded using an inverted Axio Observer microscope (Zeiss) with a Plan Apochromat N 63×/1.40 oil DIC M27 objective and a Photometrics Prime 95B camera. GFP, mCherry and JFX650 were imaged using GFP, dsRed and Cy5 filter cubes, respectively. Cells were kept at 37°C and 5% CO_2_ during imaging. Images were deconvolved using Huygens professional’s “standard deconvolution”. Line plots were generated using Fiji.

Z-stacks of HeLa cells , for colocalization analysis of FERARI subunits with SNX1, were obtained using an inverted Axio Observer microscope (Zeiss) with a Plan Apochromat N 63×/1.40 oil DIC M27 objective and a Photometrics Prime 95B camera. GFP and mCherry were imaged using GFP and dsRed filter cubes, respectively. Images were deconvolved using Huygens professional’s “standard deconvolution”. Colocalization analysis was performed in Fiji using the JaCOP plugin. Mander’s and Pearson’s coefficient were measured for 5 ROIs per cell and averaged.

HeLa WT and HeLa SNX5 + 6 KO, transfected with SNX1/Rab11/SNX5 variants were imaged using a Leica Stellaris point scanning confocal microscope with an inverted 63 x 1.4 NA oil objective. Three-minute time-series were recorded per cell with a Z-stack of 3 slices with the GFP and the mCherry channel. Transfection with different emiRFP703 constructs was verified by acquiring a stack including the emiRFP703 Channel prior to starting acquisition of the movie. Cells were kept at 37°C and 5% CO_2_ during imaging. Deconvolution was perfomed with the Leica LIGHTNING software. Kiss-and-run events were identified and quantified manually in Fiji.

### CRISPR-Cas9 KO of SNX5/6 and CRISPR-Cas9 Halo KI in the SNX1 locus

The SNX5+6 KO cell line was generated as described before (Solinger et al., 2022).

| Plasmid | Source |
| --- | --- |
| GFP-SNX1 | Kind gift from Prof. Peter J. Cullen (University of Bristol) |
| mCherry-SNX1 | Turbo GFP from Origene RG201844 was replaced with mCherry |
| mCherry-EHD1 | Kind gift from Ármín Söth from Prof. Gábor Juhász's group (Eötvös Loránd University) |
| GFP-Rabenosyn-5 | Addgene #37538 |
| Rab11FIP5-GFP | Kind gift from Prof. Rytis Prekeris (University of Colorado) |
| emiRFP703-SNX5 | Obtained by replacing GFP with emiRFP703. Original construct was a kind gift from Prof. Peter J. Cullen (University of Bristol) |
| emiRFP703-SNX5 S226E | Obtained by replacing GFP with emiRFP703. Original construct was a kind gift from Prof. Peter J. Cullen (University of Bristol) |
| emiRFP703-SNX5 F136D | Obtained by replacing GFP with emiRFP703. Original construct was a Gift from Prof. Peter J. Cullen (University of Bristol) |
| GBF1-GFP | Kind gift from Prof. Elizabeth Sztul (University of Alabama at Birmingham) |
| Arf1-GFP | Kind gift from Prof. Martin Spiess (University of Basel) |
| mCherry-Rab11 | Addgene #55124 |
| pRS315-GFP-TLG2 | Kind gift from Prof. Jeffrey Gerst (Weizmann Institute of Science) |
| pX330-U6-Chimeric_BB-CBh-hSpCas9 | Addgene #42230 |
| pBS598 EF1alpha-EGFPcre | Addgene #11923 |

### Yeast transformation

Three units of OD_600_ of yeast cells were grown in appropriate YPD or HC media to mid-log phase. Cells were spun down and washed in 1 volume of 1 × TE and 100 mM LiAc. The pellet was then resuspended in 350 μL of transformation mix (1 × TE, 100 mM LiAc, 8.5% (v/v) single-stranded DNA and 70% (v/v) PEG3350), incubated with 1 μg pRS315-GFP-TLG2 for 1 h at 42 °C, spun down (2,200 g, 2 min) and resuspended in 100 μL of HC-LEU media, and cells were plated onto selective HC-LEU media and incubated at 23 °C. This procedural description was adapted (Enkler et al., 2023).

### Live yeast imaging and FM4-64 staining

Yeast strains expressing GFP-Tlg2 were grown overnight at 23°C in selective HC-LEU medium and diluted into YPD the following morning. Cultures were grown to mid-log phase, and 1 mL of each culture was harvested by centrifugation (2,200 g, 2 min). The pellet was resuspended in 199 μL HC-complete containing 2% glucose, supplemented with 1 μL 10 mM FM4-64 (Styryl dye from Thermo Fisher Scientific (T13320)), briefly vortexed and centrifuged again (2,200 g, 2 min). The pellet was then resuspended in 200 μL HC-complete containing 2% glucose and centrifuged once more (2,200 g, 2 min) and finally resuspended in 200 μL HC-complete containing 2% glucose. Cells were imaged immediately thereafter to visualize FM4-64-labelled early endosomal compartments.

### Yeast Microscopy

Fluorescence and DIC images were acquired with an ORCA-Flash 4.0 camera (Hamamatsu) mounted on an Axio Imager.M2 fluorescence microscope with a 63x Plan-Apochromat objective (Carl Zeiss) and an X-Cite XYLIS light source with ZEN 3.9 software. Images were analysed with Fiji software. Adapted (Enkler et al., 2023).

### Image analysis using Fiji

Cells with a healthy morphology were selected using the DIC channel and the corresponding fields of views were cropped and saved for analysis. Image channels were split and background subtraction was performed on each fluorescent channel using a rolling ball radius of 10 px with sliding paraboloid. FM4-64 and GFP-Tlg2 double-positive endosomal compartments were identified, manually numbered and outlined in the GFP-Tlg2 channel using the freehand selection tool to precisely capture endosomal boundaries. Morphometric parameters were subsequently extracted from these ROIs.

### Yeast Strains

*NYY0-1 MATa ade2::ARF1::ADE2 arf1::HIS3 arf2::HIS3 ura3 lys2 trp1 his3 leu2 NYY11-1 MATa ade2::arf1-11::ADE2 arf1::HIS3 arf2::HIS3 ura3 lys2 trp1 his3 leu2 NYY18-1 MATa ade2::arf1-18::ADE2 arf1::HIS3 arf2::HIS3 ura3 lys2 trp1 his3 leu2* (Strains are from (Yahara et al., 2001)).

### Yeast two-hybrid assays

A system with two plasmids pEG202 with a DNA binding domain (DBD) and pJG4-5 (alternative name: pB42AD) with an activation domain (AD) was used. Worm genes or subdomains of the genes were cloned into the vectors using NEBuilder (New England Biolabs) cloning system to generate fusion proteins with either DBD or AD. Plasmids were then transformed into yeast strain EGY48+pSH18-34 containing a plasmid for LacZ assays. Double transformations work efficiently in yeast and resulted in strains containing all required combinations of expression vectors for experiments. 6 colonies were picked and tested for growth on plates lacking Leucine (3 are shown in the figures, but generally all 6 colonies behaved very similarly). Each growth experiment was carried out 3 times to account for differences in plates and amount of yeast streaked. Pictures were taken after 3-4 days of incubation.

### Data Availability

The data generated in this study are provided in the Supplementary Information/Source Data file.

## Acknowledgements

We would like to thank Barth Grant, Pingsheng Liu, Alicia Melendez, Peter J. Cullen, Elizabeth Sztul, Rytis Prekeris, Gabor Juhasz and Martin Spiess for *C. elegans* strains and plasmids. We are thankful to Marc Thommen and Santo Xalfa for cloning and painting some of the yeast two-hybrid assays. The IMCF of the Biozentrum is acknowledged for expert support, in particular Alexia Loynton-Ferrand and Laurent Guérard. Some *C. elegans* strains were provided by the CGC, which is funded by the NIH Office for Research Infrastructure Programs (P40 OD010440). The genomically tagged ARF-1 and SNX-1 strains were generated by Sunybiotec (China). This work was supported by the Swiss National Science Foundation (3100310_197779, 310030_219513, 320030-231859) and the University of Basel.

## Author Contribution Statement

JAS and AS conceived the study. JAS, CAF, TN, and DM performed experiments. JAS, CAF, TN, DM and AS analyzed the data. AP and FB generated the HALO-SNX1 CRISPR-KI line. JAS and AS wrote and edited the manuscript with input from all authors.

## Competing Interests Statement

The authors declare no competing interests.

---

**Supplementary Figure 1:**
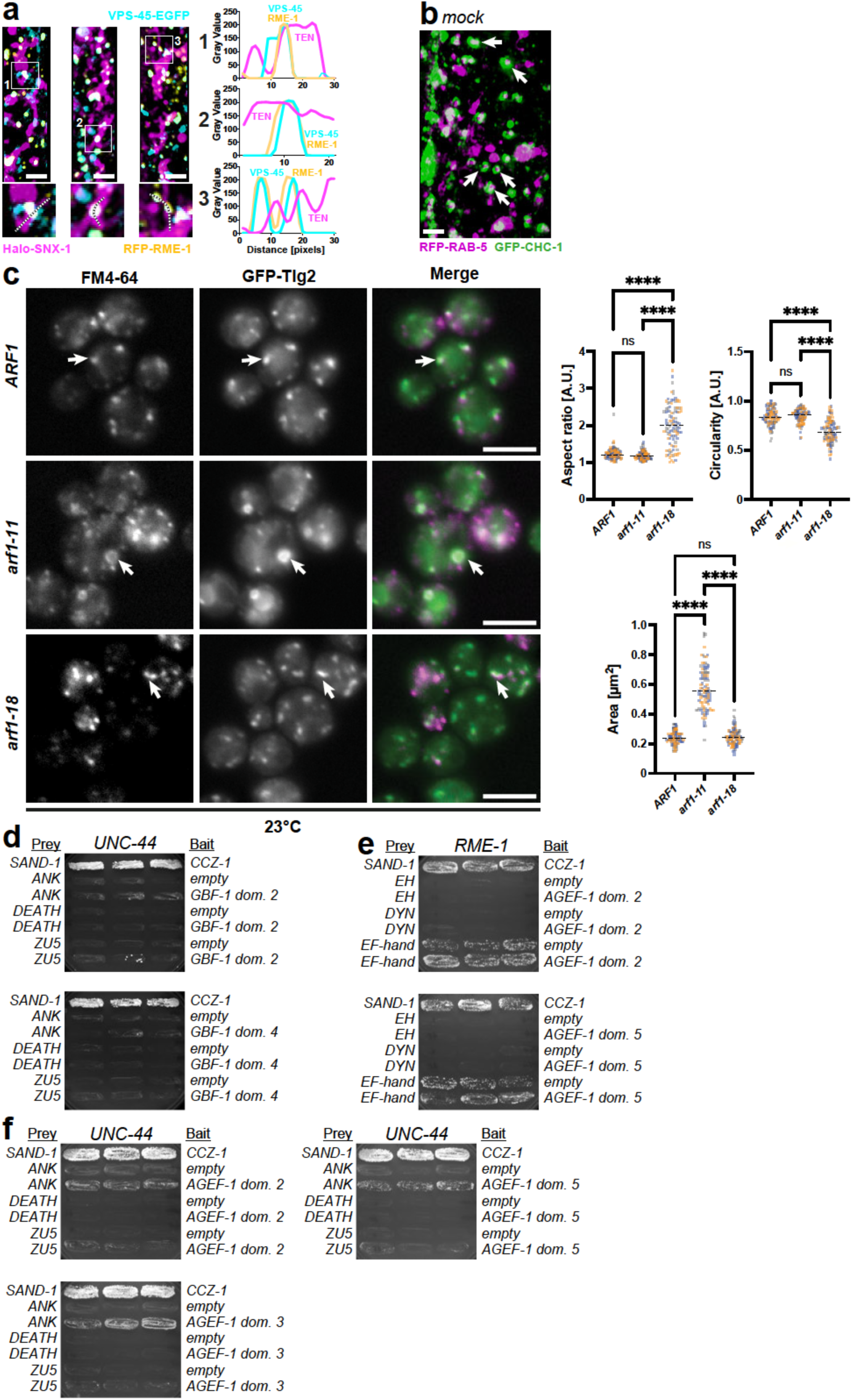
(a) 3D projections of SNX-1 tubules, closely associated with RME-1 and VPS-45 positive KAR sites. Enlargements show examples with line plots on the right. RME-1 and VPS-45 are both part of FERARI and come together on SNX-1 compartments. (b) 3D projection of early endosomes with RAB-5 and a clathrin (CHC-1) coat. These ring-shaped structures are not motile (Arrows). The moving vesicles in Fig. 1c correspond to the dot-like smaller vesicles. (c) Effects of altered Arf1 activity on the endosomal system are conserved in *S. cerevisiae*. Wild-type *ARF1*, hyperactive *arf1-11* and hypoactive *arf1-18* strains expressing the t-SNARE GFP-Tlg2 from a centromeric plasmid were stained with FM4-64 at the permissive temperature (23 °C) to label early endosomal compartments. Double-positive endosomes (GFP-Tlg2 / FM4-64) were measured in Fiji using morphometric descriptors (aspect ratio, circularity and area), followed by quantitative analysis. The hyperactive *arf1-11* allele caused enlargement of endosomes while maintaining a round shape, whereas the inactive *arf1-18* allele induced a needle-like deformation of endosomes while area remained comparable to the control. Data are shown as colour-coded individual endosomes from three biological replicates (replicate 1: 34 endosomes, grey; replicate 2: 33 endosomes, blue; replicate 3: 33 endosomes, orange; *N* = 100 endosomes per strain). Normality was not consistently met across groups, therefore nonparametric statistics were applied. Significance for aspect ratio, circularity and area were determined using Kruskal–Wallis tests followed by Dunn’s multiple comparisons. The dotted black line represents the overall median value calculated across all biological replicates. ns, not significant; ****P < 0.0001. (d) Interactions between GBF-1 domains 2 and 4 with UNC-44 domains, see Fig.3i for domains 1 and 3. (e) Yeast two-hybrid analyses of AGEF-1 domains interacting with RME-1. Complementary experiment to Fig. 4c. (f) AGEF-1 interactions with UNC-44 corresponding to Fig. 4d.

**Supplementary Figure 2:**
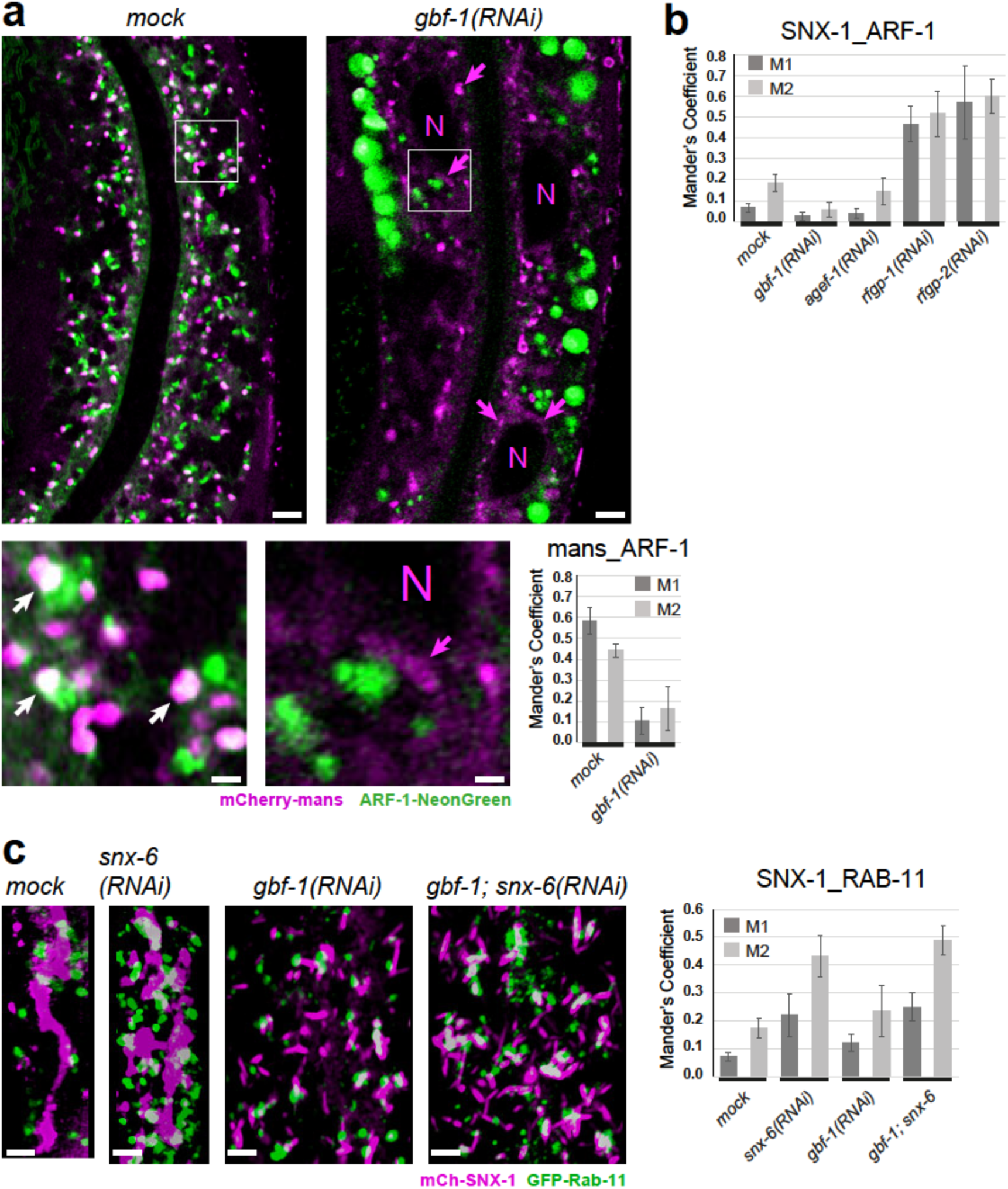
Knock-down of *gbf-1* causes Golgi disruption and some accumulation (magenta arrows) near the nucleus (N) as published before (Ackema et al., 2013). ARF-1 is frequently found on Golgi compartments (white arrows). In *gbf-1(RNAi)* ARF-1 signal is completely lost from Golgi and accumulates in large bright compartments. (b) Quantification of co-localization between ARF-1 and SNX-1 in different arfGEF and arfGAP knock-down conditions (Mander’s coefficients with average and standard deviations are shown, n=5). (c) RAB-11 accumulates in SNX-1 sorting tubules upon *snx-6(RNAi)*. It also accumulates on *gbf-1(RNAi)* induced SNX-1 needles upon *snx-6* knock-down. Mander’s coefficients are shown on the right (average and standard deviations, n=5).

**Supplementary Figure 3:**
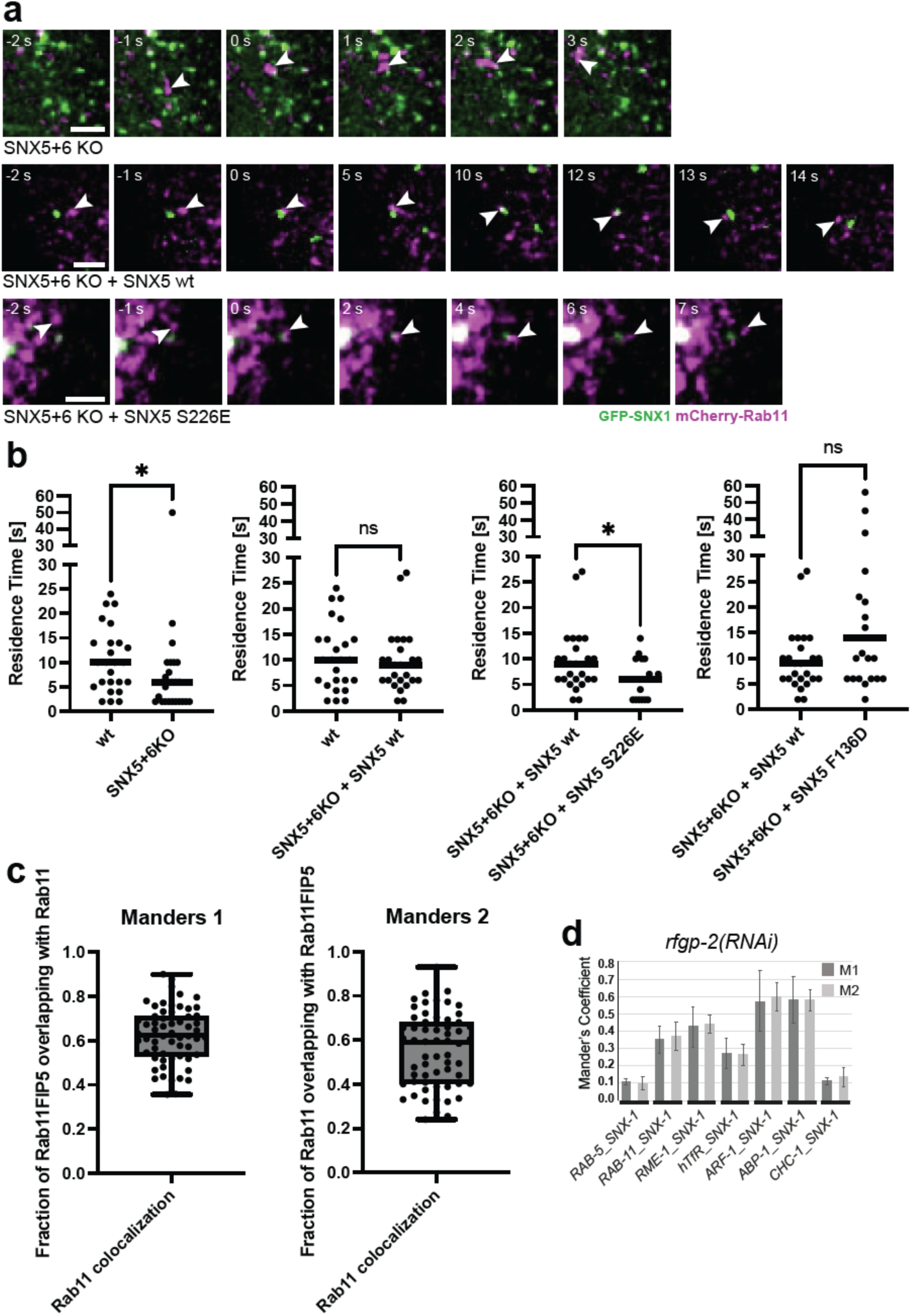
Additional kiss-and-run movie stills for Rab11 vesicles shown in Figure 6c (see Movie 5). (a) Residence times of Rab11 kiss-and-run vesicles from Figure 6d, shown as dot plots graphs. (c) Co-localization of Rab11 with Rab11FIP5 in control HeLa cells. (d) Co-localization measurements for Fig. 7a-c. Mander’s coefficients for *rfgp-2(RNAi)* condition (average and standard deviations, n=5).

## Notes

### Competing Interest Statement

The authors have declared no competing interest.

